# A data model for longitudinal multi-modal biomedical cohorts

**DOI:** 10.64898/2026.10.14.759237

**Authors:** Christian Ramirez, Ross W. Swofford, Alyssa Lima, Huong Phan, Nishigandha Phalke, Jonathan Fisher, Matthew C Lacadie, Jair Meza, Mansi Khandpekar, The Multi-Omics for Health and Disease Consortium, Zhiping Weng, Jill Moore

## Abstract

Longitudinal multimodal cohort studies require data models that can integrate heterogeneous molecular, phenotypic, and contextual data while preserving relationships across participants, biospecimens, study visits, and computational transformations. Here, we present the data model developed for the Multi-Omics for Health and Disease (MOHD) Consortium, a modality-agnostic, provenance-driven framework designed to support an evolving longitudinal biomedical resource. The model uses a compact set of linked entities and a central dataset abstraction to represent experimental, phenotypic and contextual, and computational data within a common framework. Stable identifiers, explicit provenance relationships, and flexible grouping structures enable data to be tracked across biological sources, processing workflows, and releases without requiring modality-specific restructuring of the core model. To address the complexity of participant context, we developed a harmonization framework that transforms branched REDCap survey data into a longitudinal, machine-readable representation with normalized variables and explicit response status. Interoperability with external data models and ontologies is supported through reproducible translation workflows while maintaining a stable canonical representation. We demonstrate the model through its implementation in the first production-scale MOHD data release, encompassing 956 participants, nearly 7,000 datasets, and more than 40,000 files across multiple molecular modalities and study visits. This implementation demonstrates the ability of the model to accommodate heterogeneous and longitudinal data, evolving computational products, participant withdrawal, and distinct data-access environments. Together, the MOHD implementation provides a practical framework and generalizable design principles for building longitudinal multimodal biomedical resources that remain traceable, interoperable, and adaptable as data and analytical requirements evolve.

## Introduction

Longitudinal multimodal cohort studies increasingly integrate molecular measurements with clinical, behavioral, environmental, and participant-reported information collected repeatedly from the same individuals. The scientific value of these resources depends not only on the breadth of data collected, but also on maintaining the relationships among participants, biological materials, observations, molecular measurements, and the computational transformations applied to them [1]. As studies grow across participants, timepoints, and data modalities, the ability to consistently represent, link, version, and reuse heterogeneous data becomes critical for reproducible analysis and downstream reuse [2,3].

Representing these relationships is challenging because different components of a longitudinal study operate at different biological, technical, and temporal scales. A participant may contribute multiple biological samples across study visits; the same sample may support multiple molecular assays; individual assays may generate numerous raw and processed files; and downstream computational analyses may integrate data across participants, timepoints, or modalities. Phenotypic and contextual information introduces additional complexity because some participant characteristics remain relatively stable, whereas others are measured repeatedly and change over time. Survey structure, conditional questions, and patterns of missingness must also be retained to interpret these observations correctly. A data model for longitudinal multimodal studies must therefore represent not only individual data objects, but also the biological, temporal, and computational relationships that connect them.

Existing genomic and biomedical data infrastructures provide established approaches for representing many components of this landscape. Large-scale efforts have developed data models tailored to functional genomics, including the Encyclopedia of DNA Elements (ENCODE) Project [4,5] and the Impact of Genomic Variation on Function (IGVF) Consortium [6]; rare disease studies such as the Genomics Research to Elucidate the Genetics of Rare Diseases (GREGoR) Consortium [7,8]; population-scale programs such as the All of Us Research Program [9]; and cancer-focused resources such as the National Cancer Institute Genomic Data Commons [10]. These frameworks address important challenges in experimental provenance, genomic and clinical data integration, and large-scale data sharing, but were developed around different scientific goals, data types, and operational requirements. Longitudinal multimodal cohorts bring many of these requirements together within a single analytical resource, creating a need to connect heterogeneous molecular measurements with detailed participant context while preserving their relationships over time.

The Multi-Omics for Health and Disease (MOHD) Consortium provides a practical setting in which these challenges arise simultaneously. MOHD integrates longitudinal molecular measurements with standardized phenotypic and contextual information collected across multiple disease-focused cohorts and repeated participant visits. This combination requires stable representation of participant identity, biological materials, experimental and computational data products, and longitudinal observations while accommodating heterogeneous molecular modalities and evolving analytical requirements. The scientific rationale, overall study design, and broader opportunities and challenges of the MOHD resource are described in the accompanying MOHD Comment.

Here, we describe the data model developed to address these requirements within MOHD and its implementation in a production-scale longitudinal multimodal resource. The model uses a compact set of entities and relationships to connect participants, samples, molecular and contextual datasets, computational analyses, and derived data products while retaining detailed source and processing information. We further describe a framework for harmonizing longitudinal phenotypic and contextual data, including preservation of survey logic and temporal relationships, and an approach to interoperability based on explicit translation to external standards rather than constraint of the underlying model. We demonstrate the data model through its implementation in MOHD Snapshot 1, comprising over 7,000 datasets across 956 participants, and use this production-scale implementation to identify generalizable considerations for the design of longitudinal multimodal biomedical data resources.

## MOHD Study Design

The Multi-Omics for Health and Disease (MOHD) Consortium is a longitudinal, multi-site study designed to integrate molecular measurements with detailed phenotypic and contextual information across heterogeneous disease settings. MOHD will enroll approximately 1,800 participants through six independent Disease Study Sites (DSSs), which recruit participants across different diseases, life stages, geographic settings, and genetic ancestries. Participants are followed longitudinally for up to three study visits, during which standardized biospecimens, physical measurements, and participant-reported clinical, demographic, behavioral, and environmental information are collected. The scientific rationale, cohort design, and broader opportunities enabled by this longitudinal multi-omic framework are described in the accompanying MOHD Comment.

Data generation and coordination are organized through a common consortium-wide workflow. DSSs collect biospecimens using standardized, barcoded collection kits and capture phenotypic and contextual information using harmonized REDCap instruments. Biospecimens are transferred to a centralized Omics Production Center (OPC), where genomic, epigenomic, transcriptomic, proteomic, metabolomic, lipidomic, exposomic, and metallomic measurements are generated using standardized protocols. Molecular and phenotypic/contextual data are subsequently transferred to the Data Analysis and Coordination Center (DACC), which coordinates data ingestion, accessioning, harmonization, computational processing, quality control, and release. This structure creates a common data flow from participant → biological material and contextual observations → molecular measurement → computational processing → integrated data resource, while retaining information about the site, collection event, sample, assay, and analytical processes from which each data product was derived.

### Representing biological entities and their derived data

The MOHD Data Model translates this flow of biological materials and information into a set of linked entities that preserve relationships from participants and samples through molecular measurements and derived computational data (**Fig. 1, Supplementary Table 1A**).

**Figure 1.**
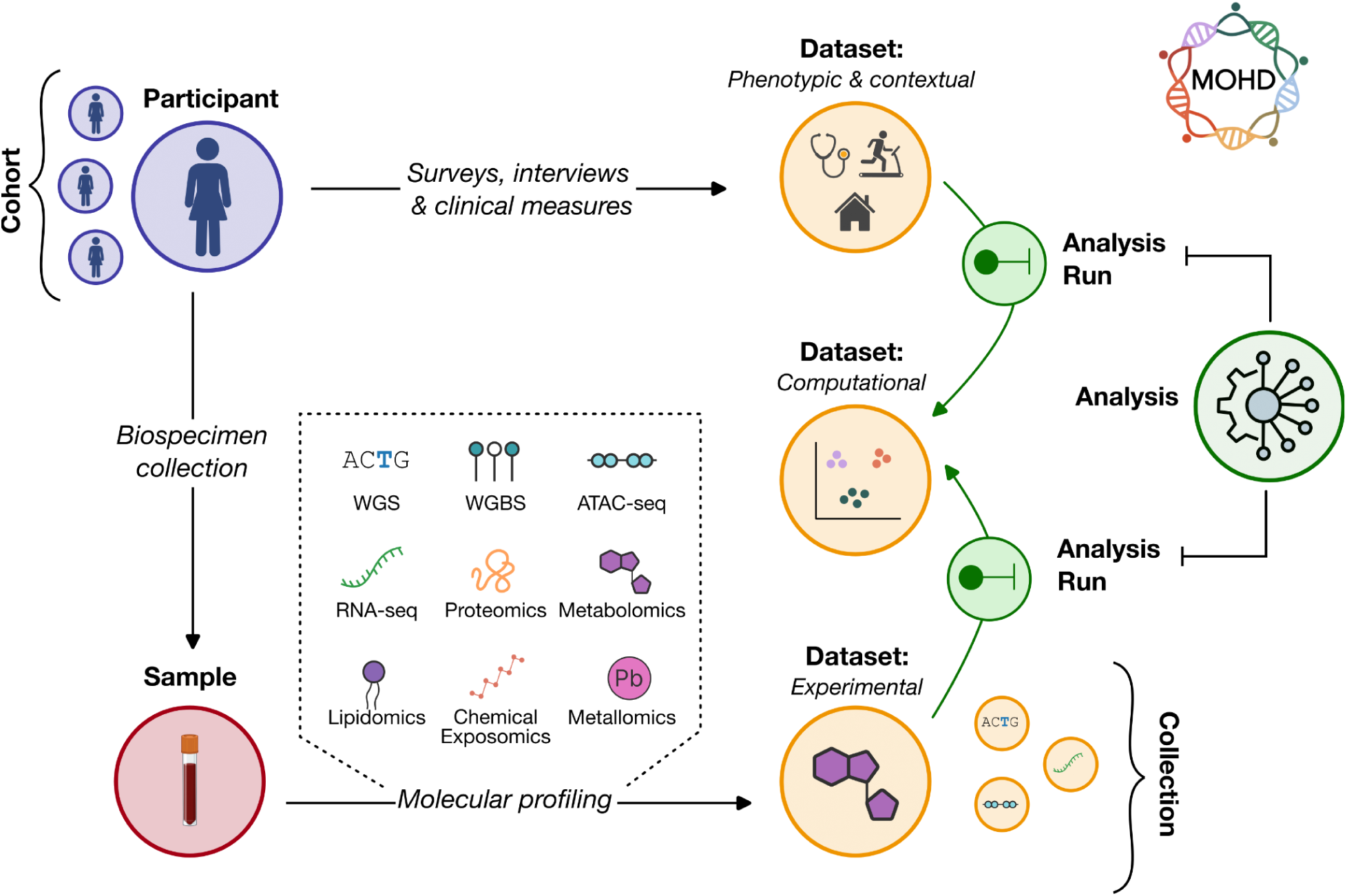
Overview of the MOHD data model. The MOHD data model organizes longitudinal, multi-omic data by linking participants, biospecimens, datasets, and computational analyses within a unified, provenance-aware framework. Participants contribute biospecimens across study visits, which are represented as samples and serve as the source for experimental datasets spanning multiple molecular modalities: whole genome sequencing (WGS), whole genome bisulfite sequencing (WGBS), ATAC-seq, RNA-seq, proteomics, metabolomics, lipidomics, chemical exposomics and metallomics). In parallel, participant-derived phenotypic and contextual data are captured as structured datasets. Computational analyses—including processing, normalization, integration, and harmonization—operate on experimental and phenotypic datasets to generate derived computational datasets. Collections group related datasets across assays or analyses, enabling coordinated data release and downstream reuse. Arrows indicate the flow of data and relationships between core components of the model.

#### Participants

Participants provide the primary longitudinal anchor for cohort data, linking observations collected across study visits to the biospecimens, molecular measurements, and phenotypic/contextual data derived from each individual. Maintaining a stable participant identity is therefore essential for integrating data generated across time, modalities, and data sources. In MOHD, each participant is assigned a globally unique, MOHD-specific identifier that remains consistent throughout data collection, processing, and release. Participant identifiers are intentionally non-informative and do not encode recruiting site or other participant attributes, allowing the same stable identifier to be used across data environments (e.g., open-access data shared through the MOHD Data Portal or controlled-access data within AnVIL). Participant attributes are instead represented as structured metadata, allowing their inclusion to be tailored to the appropriate data-access context. The participant entity contains a limited set of core attributes needed for basic filtering and cohort definition, including enrollment site, age at enrollment, sex at birth, and case or control status. More detailed clinical, demographic, behavioral, and other contextual information is retained in the harmonized REDCap datasets.

The DACC maintains the mapping between each MOHD participant identifier and the internal identifiers used by individual DSSs within REDCap. We do not require sites to adopt the MOHD accessioning scheme within their local systems, as identifier conventions may differ across sites and may support integration with existing or parallel local cohorts. Instead, sites retain their internal identifiers, while participant-level data are mapped to the common MOHD identifier during centralized ingestion and harmonization at the DACC. This approach establishes a consistent participant identity across the consortium without requiring MOHD DSSs to modify their local data-management practices.

#### Samples and biospecimen representation

Biological materials provide the physical link between participants and the molecular measurements generated from them. In the MOHD data model, samples are modeled as discrete entities representing the specific tube, aliquot, or preparation of biological material used for experimental processing. The type of biological material contained within each sample—such as whole blood, plasma, buffy coat, PBMCs, or urine—is represented as a controlled biospecimen attribute rather than as a separate entity. Representing biospecimen types using controlled vocabulary terms preserves biologically relevant information without introducing unnecessary complexity into the core data model and enables mapping to external ontologies such as UBERON. A single sample may serve as the source for multiple experimental datasets, such as whole genome sequencing and whole genome bisulfite sequencing, allowing reuse of the same biological material across assays to be represented explicitly. Each sample also retains metadata describing its collection and preparation, including collection date, processing time, tube type (e.g., Matrix, CPT, PAXgene), and the study visit or collection event associated with the sample.

Importantly, study visits are not modeled as independent entities within the core data model. Instead, temporal information is retained as metadata associated with samples and phenotypic/contextual observations. This allows the model to accommodate collections that deviate from the nominal study schedule, such as replacement or supplemental collections, without forcing them into predefined temporal categories. When analytically defined timepoints are required, such as for cross-cohort temporal analyses, observations can instead be aligned during harmonization or represented in derived computational datasets.

#### Datasets

Datasets are the central organizing unit for derived data within the MOHD data model. In the most general sense, a dataset represents a collection of files that derive from a common source; this source may be a participant, a sample, or another dataset. At present, datasets are classified into three primary categories—experimental, phenotypic/contextual, and computational—with additional dataset types incorporated as the project evolves (**Supplementary Table 1B**).

##### Experimental datasets

Experimental datasets correspond to molecular measurements generated across multiple omic modalities. Within MOHD, these include sequencing-based assays and mass spectrometry–based measurements spanning ten distinct modalities. The source of each experimental dataset is a specific sample. Because participants contribute multiple samples across study visits and biospecimen types, modeling experimental data at the sample level retains information that would be lost if molecular measurements were linked only to participants. This granularity also supports troubleshooting and quality control by allowing issues with individual collection tubes, sample handling or storage, and potential sample swaps to be traced to the specific material used for an assay.

Files associated with experimental datasets include raw data generated directly by the assay, such as FASTQ files for sequencing-based experiments or mzML files for mass spectrometry. Experimental datasets may also include primary processed outputs derived from these data, such as aligned reads, peak calls, or quantification files. In contrast, downstream products derived from multiple experiments or samples are not included within individual experimental datasets. These products are instead modeled as separate computational datasets, as described below.

##### Phenotypic/contextual datasets

Phenotypic and contextual information is often represented as participant-level attributes or as a flat matrix in which each row corresponds to a participant and each column to a measured variable. This representation becomes difficult to maintain in longitudinal studies, where the same information may be collected repeatedly over time, and in highly structured surveys where the interpretation of individual values depends on instrument structure, branching logic, and collection context. Rather than modeling these observations as attributes of the participant entity, the MOHD data model represents phenotypic/contextual information as datasets derived from participants.

Treating phenotypic/contextual information as datasets allows the same underlying observations to support multiple representations without altering the participant entity or the source data. These include source REDCap exports provided by the DSSs, harmonized longitudinal representations generated by the DACC, and derived flat tables created for specific analytical uses. These datasets can be versioned and processed using the same mechanisms applied to molecular data, allowing transformations such as harmonization or derivation of analysis-ready variables to be represented through explicit computational workflows. The approach also preserves the original longitudinal and survey structure while allowing simplified representations to be generated when appropriate. The transformation of heterogeneous REDCap data into a common longitudinal representation is described in detail below (**Figure 2**).

**Figure 2.**
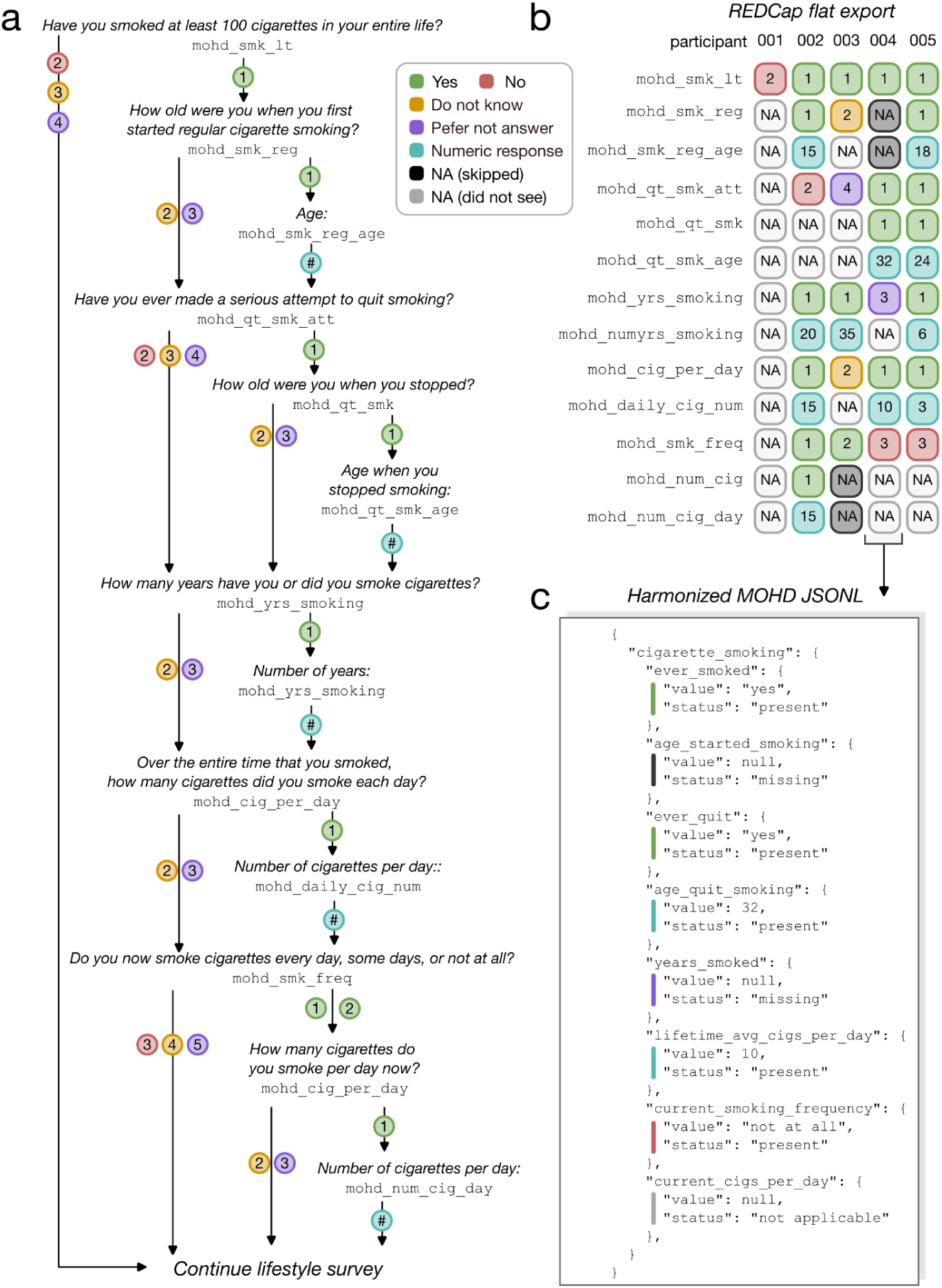
Harmonization of branched REDCap survey data into a structured, machine-readable representation. **a,** Example branching logic for smoking-related questions from the MOHD core REDCap survey. Conditional logic determines which follow-up questions are displayed based on prior responses, resulting in many questions being unseen or inapplicable for a given participant. Colored circles represent possible responses in the REDCap instrument, including categorical responses and numeric values. **b,** Illustration of how these responses appear in a standard REDCap CSV export across a set of five mock participants. The flat tabular structure does not encode branching logic or question dependencies, making it difficult to distinguish between values that are missing because a question was not shown, intentionally skipped, or not applicable. Boxes, representing observed responses from mock participants, are colored as in **a**, with two levels of NA corresponding to intentionally skipped by the participant (dark gray) or not applicable due to branching logic (light gray). **c,** Corresponding harmonized MOHD JSONL representation for mock participant 004. Each survey variable is encoded with a normalized value and explicit status field (e.g., present, missing, not applicable), while the original REDCap export is retained as the authoritative source record.

##### Computational datasets

The MOHD data model also supports computational datasets generated through consortium-wide analyses and annotation efforts. These include harmonized and batch-corrected matrices for molecular assays, annotations of cis-regulatory elements, and outputs from machine learning and deep learning models. Computational datasets may be derived from individual experimental or phenotypic/contextual datasets or from groups of datasets termed collections (discussed below). By modeling computational products as datasets with defined sources and associated files, derived products can be managed using the same framework as primary measurements while retaining their relationships to the data and analyses from which they were generated.

#### Grouping data across biological, technical, and release contexts

##### Cohorts

Cohorts represent defined groups of participants and are used to capture biologically or clinically meaningful subsets of the study population. Cohorts within MOHD include participants recruited by a specific DSS, as well as broader groupings defined by study design, enrollment period or participant attributes (e.g., age, sex, or disease status). Cohorts are used as inputs to analyses that aggregate phenotypic/contextual data across participants, such as the harmonized REDCap representations, and to define participant groups associated with specific study phases or releases. For example, Snapshot 1 includes six DSS-specific cohorts and a release-level cohort containing all participants included in the data freeze.

##### Collections

Collections are groupings of datasets and provide a flexible mechanism for organizing data across assays, timepoints, and analytical contexts. They may include datasets from a single modality or span multiple data types. Collections can represent groups of experimental datasets generated under shared technical conditions, such as those produced within the same sequencing or mass spectrometry run (i.e., a batch). This preserves information relevant to quality control, batch effect assessment, and harmonization without requiring a separate batch entity. Collections may also represent inputs to computational analyses, such as joint genotyping of whole genome sequencing datasets or generation of harmonized proteomics quantifications. At the release level, a collection can similarly define the complete set of datasets included in a data freeze, complementing the cohort that defines its participant membership. Using a single grouping construct for these purposes avoids introducing separate entity types for each use case, while the purpose of each collection is specified through its associated metadata.

#### Representing computational processing and provenance

##### Analysis and analysis runs

To distinguish computational methods from their individual executions, analyses and analysis runs are modeled as distinct entities within the MOHD data model. An analysis represents a computational pipeline, workflow, notebook, or script that can be applied to one or more datasets. An analysis run corresponds to a single execution of an analysis, capturing the precise context under which it was performed. For example, the MOHD RNA-seq uniform processing pipeline is accessioned as a single analysis entity (MOHD_AP100002), while each execution of this pipeline on an individual RNA-seq dataset is represented as a distinct analysis run. For each analysis run, the data model records the analysis definition used, including its accession and version, together with a single input and output accession, each of which may correspond to either a dataset or a collection. Collections enable analyses to consume or generate multiple datasets while retaining this simple input–output structure and preserving the identity of the individual datasets involved.

##### Files as products within datasets

Computational processing does not necessarily result in the creation of a new dataset. When processing produces files that remain part of the same underlying data product, raw and processed files are retained within the originating dataset. For example, an RNA-seq dataset may contain the raw FASTQ files, BAM files of aligned reads, and TSV gene quantification files generated by the MOHD uniform RNA-seq processing pipeline. The corresponding analysis run therefore has the same dataset as both its input and output, while recording the computational transformation that generated the additional files. This approach avoids creating a new dataset for every intermediate stage of processing, which would substantially increase the complexity of the data model and fragment a conceptually unified data product across multiple entities. New computational datasets are instead created when processing produces a distinct derived data product, such as outputs from secondary analyses beyond uniform primary processing, particularly when those outputs integrate information across multiple datasets or samples. This distinction allows datasets to remain stable organizing units while analysis runs capture the computational transformations applied to them.

##### Manifests and file-level lineage

Because multiple analysis runs may contribute files to the same dataset, such as through processing against different genome reference versions, provenance is also tracked at the file level. Each analysis run includes input and output manifests that list the files consumed and generated during that execution, linking individual files to the computational process responsible for their generation. File records additionally include checksums and other identifying metadata to support integrity validation and reproducible tracking across storage and analysis environments.

Together, dataset-level relationships and file-level manifests provide complementary layers of provenance. Dataset relationships describe the broader lineage among biological sources and derived data products, while analysis runs and manifests record the specific computational steps through which individual files were generated. This allows detailed processing histories to be reconstructed without requiring every intermediate file or processing step to become an independent dataset.

### Stable identity across an evolving resource

Stable identifiers allow data and their relationships to remain consistently referenced as metadata, analyses, and data releases evolve. To support this, we developed a standardized accessioning scheme for entities in the MOHD data model, with deliberate exceptions for files and samples described below. Most entities are assigned stable identifiers that uniquely distinguish them within the system while encoding a limited set of intentionally selected information. The scheme balances uniqueness and human interpretability without attempting to encode the full biological, technical, or participant metadata associated with an entity.

#### General accessioning principles

All non-sample accessions begin with the prefix MOHD, indicating that the entity originates from the MOHD project (**Supplementary Table 1C**). This project-level namespace allows MOHD entities to remain identifiable when integrated with other resources and distinguishes data products generated as part of the MOHD project from those generated through related efforts. For example, DSSs may generate additional datasets from the same participants through ancillary studies or externally funded projects. The data model can accommodate these datasets while assigning them distinct project-level namespaces that preserve their provenance.

Following the project prefix, each accession includes a two-letter code denoting the entity type. Distinct codes are defined for experimental datasets (e.g., whole genome sequencing, RNA-seq), computational datasets, collections, participants, cohorts, analyses, and analysis runs. The coding scheme is intentionally extensible: new entity or dataset types can be incorporated by defining additional two-letter codes without modifying existing accessions. These codes provide lightweight, human-interpretable cues that allow users to distinguish broad entity and dataset types when working in environments where associated metadata may not be immediately visible, such as remote servers or cloud storage. The codes are not intended to encode a complete classification of the entity; detailed attributes are instead maintained within the entity metadata. This distinction preserves practical interpretability without requiring the accessioning scheme to anticipate or encode every possible dataset subtype. Additionally, because participant and contextual attributes are maintained separately from the identifier itself, the same stable accession can also be used across controlled- and open-access environments while varying the metadata exposed in each context.

Each code is followed by a six-digit numerical identifier. For the current phase of MOHD, identifiers begin with the digit “1,” providing a mechanism to distinguish entities generated during future study phases if needed. Within this identifier space, numerical identifiers are assigned sequentially to support internal data coordination and tracking. However, their ordering is not intended to represent the timing of biological collection, data generation, or processing, which may occur asynchronously across sites and modalities. Temporal relationships are therefore represented through explicit metadata and harmonized data products rather than inferred from accession numbers. The identifier space is sufficiently large for anticipated entities and can be expanded for high-volume entity types, such as analysis runs, by extending the range of leading digits if needed.

#### Practical exceptions

The accessioning scheme described above provides a useful default, but it is not applied uniformly when an alternative identifier better supports data use or provenance. We make two such exceptions. Files use descriptive identifiers that help users recognize their contents, whereas samples retain identifiers established during physical biospecimen collection. These exceptions preserve practical information that would be lost by assigning fully independent, pseudo-random accessions.

##### Files

File identifiers derive from their associated dataset accessions rather than being fully independent (**Supplementary Table 1D**). Each file identifier incorporates the parent dataset accession together with a curated suffix describing the file’s contents, key attributes such as genome build or annotation version, and, where relevant, the version of the processing pipeline used to generate it. This structure preserves the relationship between a file and its parent dataset while allowing users to distinguish commonly used files without repeatedly consulting external metadata.

For example, the MOHD RNA-seq pipeline generates multiple output files, including distinct BAM files containing reads aligned to the genome and transcriptome, respectively. With a fully randomized file-naming scheme, these files would be difficult to distinguish without cross-referencing metadata tables. Instead, standardized descriptors are appended to the dataset accession; for example:

MOHD_ER100001_align_GRCh38_v0 and
MOHD_ER100001_align_GENCODE49_v0

allow users and computational workflows to distinguish the relevant BAM files directly from their identifiers. To ensure consistency across the data model, these suffixes are drawn from controlled, assay-specific vocabularies maintained by the DACC rather than entered as free text.

Index files required by many genomic tools (e.g., BAM/CRAM or VCF indexes) present a separate practical consideration. Assigning these files independent identifiers can break the matching filename stems expected by standard tools, whereas omitting them from the model requires users to regenerate them before analysis. MOHD therefore treats index files as deterministic sidecar files: they retain the exact filename stem of their corresponding primary file (e.g., MOHD_ER100001_align_GRCh38_v0.bam.bai) and are recorded in analysis-run output manifests rather than accessioned as independent data objects. Checksums and provenance information remain associated with these files, allowing them to be tracked without compromising compatibility with existing genomic software.

##### Samples

Samples represent a second intentional exception because their identifiers originate during the biospecimen collection and tracking process (**Supplementary Table 1E**). MOHD uses standardized sample collection kits assembled centrally and distributed to DSSs, with barcoded tubes tracked throughout collection, processing, and shipping. Sample identifiers encode the originating DSS using a three-letter code followed by a four-digit number, together with a tube-specific suffix indicating biospecimen type and tube number. Rather than replacing these operational identifiers with independently generated MOHD accessions, the original sample identifiers are retained within the data model. This maintains continuity between the physical material collected at the DSS, its processing and shipment records, and the molecular datasets ultimately generated from that sample, while avoiding an additional identifier-mapping layer that could introduce ambiguity or error. When the same sample is used for multiple assays—for example, whole genome sequencing and whole genome bisulfite sequencing—the shared sample identifier provides a direct means of identifying datasets generated from the same physical source.

### Harmonizing longitudinal phenotypic and contextual data

Phenotypic and contextual data present a distinct harmonization challenge in longitudinal, multi-modal studies. The MOHD consortium developed a core REDCap survey to uniformly capture information across a wide range of domains, including clinical and medical history, social and structural determinants of health, and lifestyle and health behaviors for all MOHD participants. In total, the core survey comprises more than 700 distinct variables (**Supplementary Table 2**) collected across six independent DSSs and multiple study visits. In addition to this common core, individual sites may collect disease- or age-relevant information through additional instruments, resulting in thousands of unique variables at some DSSs.

Although the consortium has standardized the core REDCap instruments, the resulting data are inherently heterogeneous. For example, extensive branching logic further determines which questions are presented to individual participants. For example, participants who have never smoked are not shown follow-up questions about smoking behaviors (**Figure 2a**), while questions related to pregnancy or menopause are presented only to adult participants recorded as female at birth. As a result, only 280 questions are expected to be presented to all MOHD participants, with more than 500 variables conditionally displayed based on participant characteristics or prior responses. An empty value in the resulting data therefore does not have a single interpretation: it may indicate that a question was shown but left unanswered, that it was not applicable and therefore never displayed, or that the relevant information was not collected during that study event.

Longitudinal collection introduces an additional dimension. Some participant characteristics, such as race and ethnicity, are generally collected once and remain applicable across the study, whereas other contextual measures, including diet, sleep patterns, health behaviors, and clinical status, may change over time and are therefore collected repeatedly. Consequently, harmonization must preserve both stable participant attributes and time-varying observations while retaining the study event at which each response was collected.

#### Limitations of flat exports

To protect participant privacy and minimize the movement of personally identifiable information, MOHD was designed so that each DSS maintains and controls access to its own REDCap system. This approach keeps identifiable source data localized at the recruiting sites and limits its movement across the consortium. As a result, the DACC does not have direct access to site-specific REDCap systems, and phenotypic and contextual data are instead transferred through periodic standardized exports rather than direct database integration. The default REDCap export format is a flat CSV in which survey structure, branching dependencies, response definitions, and longitudinal relationships must be interpreted using the accompanying codebook and knowledge of the underlying instruments. As the number of participants, study visits, and variables increases, reconstructing these relationships independently for each analysis becomes increasingly impractical.

Flat exports also reflect the structure in which data were collected rather than necessarily the structure most useful for cross-participant or cross-cohort analysis. For example, a participant may need to come in for a supplemental biospecimen collection if they were unable to give blood during their first enrollment visit. This sample would need to be associated analytically with a participant’s baseline timepoint, while the original collection event remains important for provenance. We therefore distinguish the source representation, which preserves when and how information was collected, from the harmonized representation, which can establish consistent longitudinal relationships for downstream use.

#### A common harmonized representation

To provide a consistent representation across sites and study visits, the DACC developed a harmonization framework that transforms standardized REDCap exports into a longitudinal, machine-readable JSONL representation. Rather than treating each export as an independent flat table, the harmonized structure organizes observations hierarchically by participant and study event, with individual responses organized by survey domain and accompanied by their value and status. The original REDCap exports are retained as the authoritative source records, allowing the harmonized representation to prioritize interpretability and downstream use without sacrificing provenance.

Conceptually, the representation follows the hierarchy:

participant → study event → survey domain/instrument → observation

Within this structure, REDCap variable names are replaced with standardized, human-readable identifiers that provide greater semantic context while remaining suitable for programmatic use. For example, the question “Have you smoked at least 100 cigarettes in your entire life?” is represented in the source REDCap export as mohd_smk_lt, with numeric response codes corresponding to “Yes,” “No,” “Don’t know,” and “Prefer not to answer” (**Fig. 2B**). In the harmonized representation, this observation is named ever_smoked, placed within the hierarchy lifestyle → substance_use → tobacco_use → cigarette_smoking, and its numeric response is translated to the corresponding human-readable value (**Fig. 2C**). This organization reduces the need for users to repeatedly consult REDCap codebooks to interpret variable names and responses.

Each observation also includes a status field that distinguishes whether a response is present, unanswered, or not applicable (**Fig. 2C**). For example, if a participant reports never smoking, conditional downstream questions about smoking behavior are represented as not applicable, whereas a question that was presented but left unanswered can be identified as missing. Preserving this distinction prevents structurally absent responses from being conflated with conventional missing data and allows downstream computational workflows to interpret response status without reconstructing the original REDCap branching logic.

This harmonized structure also allows observations to accumulate across study visits without replacing information collected previously. The enrollment visit generally contains the most comprehensive set of phenotypic and contextual data, including relatively stable participant characteristics, whereas subsequent visits selectively repeat measures expected to change over time, such as diet, sleep, health behaviors, and clinical characteristics. Each observation remains associated with the study event at which it was collected, allowing changes over time to be examined while retaining the complete history of previously collected information.

### Translation to external data models and ontologies

To support interoperability with existing biomedical resources while preserving the full structure of the MOHD data model, the DACC develops explicit translation workflows that map harmonized MOHD representations into external data models and ontologies. Interoperability is particularly important for longitudinal multi-modal resources because individual data domains may need to interface with different community standards, and those standards may evolve over the lifetime of the study. Maintaining a stable canonical representation while translating to external standards allows these requirements to be accommodated without repeatedly restructuring the underlying data resource. Rather than constraining the canonical MOHD representation to conform to a single external standard, these workflows generate task-specific representations while preserving MOHD as the authoritative source. For example, participant-level phenotypic data can be translated into formats compatible with the OMOP Common Data Model, while biospecimen types can be mapped to anatomical ontologies such as UBERON.

Translation workflows are implemented as reproducible notebooks and scripts and are made available to researchers through the MOHD AnVIL workspaces. As an initial use case, we provide an example workflow translating smoking-related survey variables from the MOHD harmonized JSONL format into the OMOP common data model, illustrating how MOHD encodes response values alongside explicit status fields (e.g., present, missing, not applicable) and how these distinctions are preserved during translation. Additional translation workflows may be developed in response to community needs. Users may request guidance or propose new translations through the MOHD Data Portal, and the DACC provides documentation, example notebooks, and hands-on training (e.g., workshops or tutorials) to support community-led development. In cases of sustained demand or broad utility, the DACC may formally curate, version, and release an official translation workflow as part of the MOHD data resource.

### Application to a production-scale longitudinal multi-omic release

We evaluated the data model through its implementation in MOHD Snapshot 1, the first production-scale MOHD data release (**Fig. 3**, **Table 1**). Snapshot 1 includes data from 956 participants recruited across six DSSs, encompassing nearly 7,000 datasets and more than 40,000 files across multiple molecular modalities, phenotypic and contextual data, and derived computational products. Snapshot 1 provided a practical test of whether the data model could accommodate heterogeneous data generated across independent recruiting sites, molecular technologies, biological materials, and computational workflows without requiring modality-specific restructuring of the core model. The release also tested the longitudinal aspects of the model. Although most participants in Snapshot 1 are represented primarily by baseline data, participants from cohorts with intentionally compressed study timelines, such as those in the pre-eclampsia cohort, contributed data from later study visits, including some participants with observations across all three planned timepoints. These later observations were incorporated alongside baseline data without requiring modification of the underlying data model.

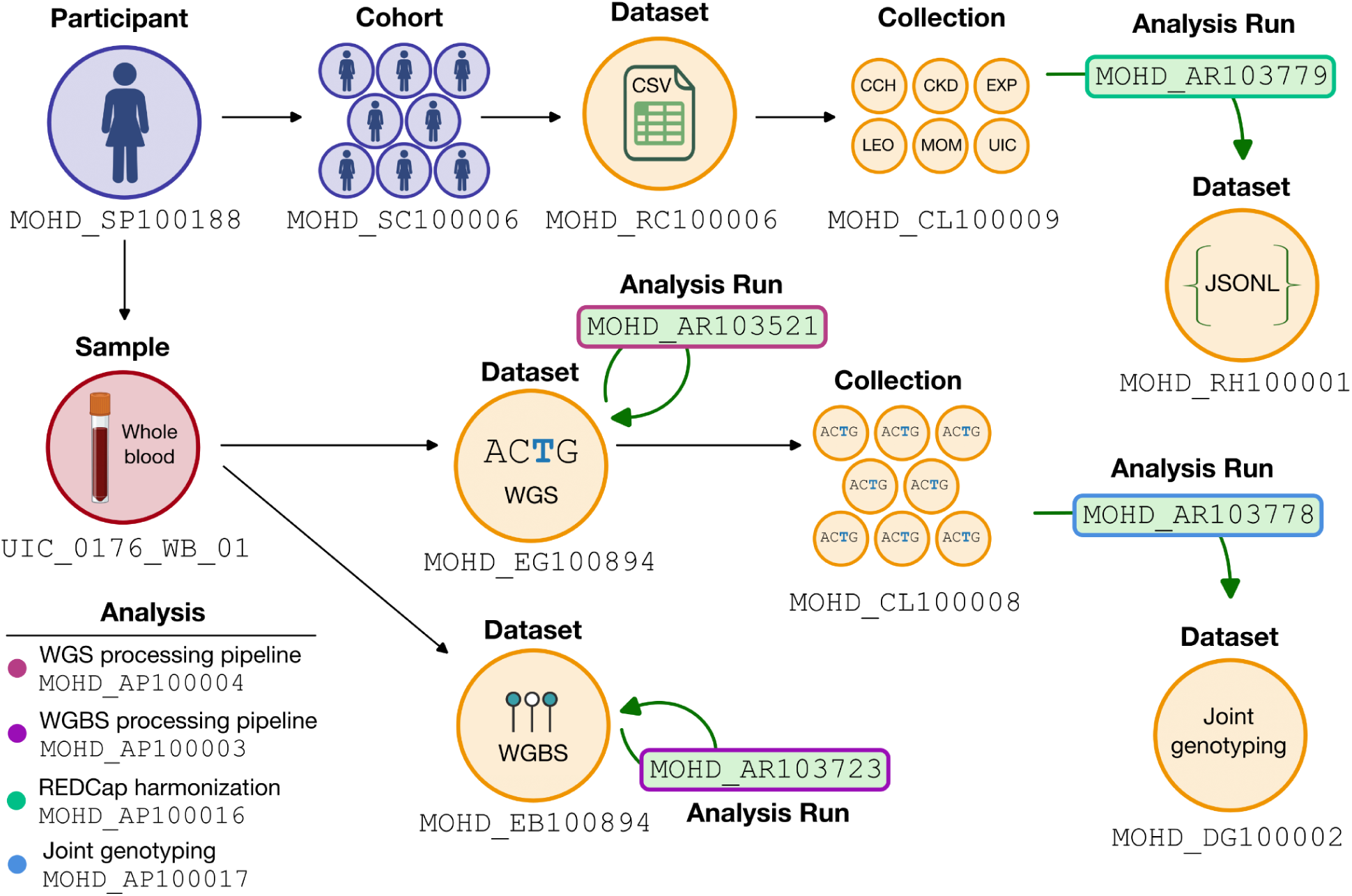

**Table 1.** Summary of Snapshot #1.

| Category | # datasets | # files | # analysis runs |
| --- | --- | --- | --- |
| WGS | 915 | 6,433 | 915 |
| WGBS | 915 | 12,810 | 915 |
| ATAC-seq | 492 | 4,920 | 492 |
| RNA-seq | 922 | 11,064 | 922 |
| Metabolomics | 1,210 | 3,577 | 1 |
| Lipidomics | 1,211 | 2,372 | 1 |
| Chemical exposomics | 690 | 2,679 | 3 |
| Metallomics | 612 | 612 | 2 |
| Aggregate analysis | 16 | 64 | 9 |
| **Total** | 6,983 | 44,531 | 3,260 |

Snapshot 1 additionally provided an opportunity to test whether the model could accommodate changes in participant data-sharing status after data had already entered the resource. In one case, a participant withdrew from the study and requested destruction of previously collected data. Because datasets, samples, files, and computational products remain explicitly linked to stable participant and source identifiers, affected data products could be systematically identified and removed while preserving the integrity of the remainder of the release. Although participant withdrawal is an expected consideration for longitudinal cohort studies, explicit relationships among participants and their derived data products facilitate systematic identification of data affected by changes in consent or sharing status.

Finally, Snapshot 1 tested whether the same data model could support distinct data-access environments with different users and technical requirements. Within AnVIL, participant-level data are accessed primarily by computational researchers, and the MOHD data model must integrate with the entity structures and technical requirements of the Terra/AnVIL environment. In contrast, the MOHD Data Portal is designed to support a broader range of users through interactive visualization, exploration, and point-and-click access to open data products, placing greater emphasis on human-interpretable identifiers and metadata. The combination of stable accessions, structured metadata, and a common underlying representation allowed Snapshot 1 to be implemented across both environments without maintaining separate data models for each. Thus, the same model could satisfy the computational and technical integration requirements of AnVIL while supporting the accessibility and interpretability required by the MOHD Data Portal.

### Generalizable principles for longitudinal multimodal data architecture

Several general design principles emerged through iterative development of the MOHD data architecture and its implementation in a production-scale longitudinal multi-omic resource. Together, these principles reflect practical considerations for balancing simplicity, provenance, usability, and interoperability as longitudinal biomedical resources evolve.

#### Keep the core data model simple by assigning identity only where it is needed

Not every concept in a complex longitudinal study requires an independent entity or accession. We found it useful to reserve persistent identity for objects that must be independently referenced or tracked over time, such as participants, samples, datasets, analyses, and analysis runs, while representing other concepts through structured metadata or flexible relationships. For example, biospecimen type and study visit are represented as attributes, whereas collections can flexibly capture technical batches, analytical groupings, or data releases without introducing a new entity type for each use case. These choices should reflect the underlying study design: MOHD uses a relatively constrained set of standardized biospecimen types, disease states, and molecular assays, whereas studies with greater heterogeneity in these dimensions may benefit from representing additional concepts as independent entities. Thus, complexity in the underlying study does not necessarily require equivalent complexity in the core data model, but the appropriate level of abstraction will vary across resources.

### 3Treat phenotypic and contextual data as a first-class data modality

Phenotypic and contextual information is often represented as a collection of tables associated with molecular datasets, but longitudinal cohorts can require a richer representation. In MOHD, these data are accessioned as datasets within the same core model used for molecular and computational data, allowing them to retain stable identity, provenance, and explicit relationships to participants and study events. This approach is particularly valuable when participant context includes complex survey structure, repeated observations, conditional relationships among variables, and information collected across multiple sites. Treating participant context as a first-class data modality allows its structure and longitudinal relationships to be harmonized with the same rigor applied to molecular measurements, rather than relegating it to auxiliary metadata.

#### Preserve source information while allowing analytical representations to evolve

Longitudinal studies may operate for many years, during which reference files, software versions, processing methods, and analytical requirements will inevitably change. We therefore preserve the most primary form of the source data while allowing multiple processed or harmonized representations to be generated and explicitly linked back to it. This principle applies equally to experimental and phenotypic/contextual data: raw molecular measurements can be reprocessed using updated computational methods, while original REDCap exports remain available as the authoritative source for evolving harmonized representations. Designing for this evolution from the outset allows analytical representations to change without overwriting previous products or obscuring the source data from which they were derived.

#### Design for both human and machine use

Longitudinal biomedical resources must support users with different levels of computational expertise and across multiple data-access environments. Data representations should therefore balance human interpretability with the standardization and structure required for programmatic analysis, allowing researchers to understand and navigate the data while enabling computational workflows to reliably identify entities, relationships, and metadata. Machine-actionable representations will become increasingly important as machine learning, multimodal modeling, and AI-assisted analytical approaches become more widely used in biomedical research. Designing for both forms of interaction from the outset allows the same underlying resource to support interactive exploration and conventional analysis while remaining suitable for increasingly automated computational workflows.

#### Design for interoperability without constraining the canonical model

Biomedical data resources rarely operate in isolation and may need to interface with multiple external data models, ontologies, and analytical platforms over their lifetime. Rather than designing the canonical representation around a single external standard, we found it useful to maintain a stable internal model and support interoperability through explicit, reproducible translations. This approach allows different data domains to adopt appropriate community standards while insulating the underlying resource from changes in those standards. Interoperability can therefore be treated as an extensible layer around the data model rather than a constraint that determines its internal structure.

### Conclusion

Longitudinal multimodal studies require data models that preserve the relationships among participants, biological materials, molecular measurements, contextual observations, and computational transformations as data accumulate across time. The MOHD data model provides one implementation of this approach, using a compact set of linked entities, stable identifiers, explicit provenance, and harmonized participant context to support an evolving multi-omic resource. Its application to a production-scale MOHD data release demonstrates that these principles can accommodate heterogeneous data types and longitudinal observations while maintaining traceability and interoperability. The specific implementation reflects the scientific and operational structure of MOHD and is not intended as a universal schema; studies with different levels of standardization, decentralization, or data heterogeneity may require different entities and relationships. More broadly, the design considerations described here provide an adaptable framework for developing data resources that remain interpretable, reusable, and extensible as longitudinal biomedical studies grow in scale and complexity.

## Author Contributions

J.E.M. conceived and designed the data model, led data harmonization, data management, and data access framework development, and wrote the manuscript. C.R. developed the data processing pipelines and performed initial data processing. H.P. contributed to data processing. R.W.S. and A.L. coordinated data integration and management across the consortium, and A.L. developed the harmonized REDCap instruments. N.P., J.F., M.C.L., J.M. and M.K. developed and designed the MOHD Data Portal. J.E.M. and Z.W. provided overall project leadership and secured funding. The Multi-Omics for Health and Disease Consortium contributed to evaluation of the data model design, provided feedback on feasibility and implementation, and reviewed and approved the manuscript. All authors contributed to manuscript editing and approved the final version.

## Acknowledgements

This study was funded by the National Institutes of Health grant U01HG013198 (Z. W.). We thank the individuals who participated in the MOHD study for their invaluable contributions to this research.

## Data Availability

All MOHD data are consented for general research use and are available through the NHGRI Analysis, Visualization, and Informatics Lab-space (AnVIL) via controlled access in dbGaP under accession phs003765.v1.p1. Access to individual-level genomic and associated molecular data is available to qualified researchers through dbGaP in accordance with NIH data access policies. Summary-level data, metadata, and additional resources will be accessible through the MOHD Data Portal (https://data.mohdproject.org).

### Software availability and reproducibility

All data ingestion, accessioning, and harmonization workflows are implemented as version-controlled scripts maintained by the MOHD DACC. Source code for these workflows, including REDCap harmonization, molecular data ingestion, and accession snapshotting, is publicly available (https://github.com/MOHD-Consortium) and versioned using tagged releases. Each data freeze is associated with a specific software version to ensure reproducibility and traceability across releases.

## The MOHD Consortium Authorship

### Disease Study Site Awards (contact PI, MPIs, Co-Is, other members, all alphabetical by last name)

#### U01CA288325

Joseph B. McCormick^1,2^, Jennifer E. Below^3,4^, Kari E. North^1,2^, Susan P. Fisher-Hoch^1,2^, Mariaelisa Graff^2^, Absalon D. Gutierrez^5^, Heather M. Highland^1,2^, Miryoung Lee^1,2^, Kristin L. Young^1,2^, Ana L. Crow^1,2^, Elena F. Evans^3,4^, Elizabeth G. Frankel^3,4^, Joshua M. Landman^3,4^, Anna S. Lorenz^3,4^, Marcela Morris^1,2^, Autumn R. Morrow^3,4^, Norma P. Perez-Olazaran^1,2^, Alexander Petty^3,4^, Rashedeh Roshani^3,4^, Ravi Shah^6^, Rocio Uribe^1,2,7^, Ting-Chen Wang^3,4^, Wanying Zhu^3,4^

1. Department of Epidemiology, Border Health Research Center, School of Public Health, Brownsville Regional Campus, The University of Texas Health Science Center at Houston, USA

2. Department of Epidemiology, Gillings School of Global Public Health, The University of North Carolina at Chapel Hill, USA

3. Vanderbilt Genetics Institute, Vanderbilt University Medical Center, USA

4. Department of Medicine, Division of Genetic Medicine and Clinical Pharmacology, Vanderbilt University Medical Center, USA

5. Department of Medicine, Division of Endocrinology, Diabetes, and Metabolism, The University of Texas Health Science Center at Houston, USA

6. Vanderbilt Diabetes Center, Vanderbilt University Medical Center, Nashville, TN

7. The Multi-omics for Health and Disease Consortium Community Partner Panel Member

#### U01HG013201

Krzysztof Kiryluk^8^, George Hripcsak^9^, Iuliana Ionita-Laza^10^, Gary W. Miller^11^, Ana Navas-Acien^11^, Maya Sabatello^12^, Ning Shang^8^, Shayan Shirazian^8^, Teena Zachariah^8^, Estefhany D. Disla^8^, Oleksandr Kravets^8^, Lucia Martinez^8^, Jessica R. Singer^13^, German Varela^8^, Sebastian Villegas^8^, Chen Wang^8^

8. Division of Nephrology, Department of Medicine, Columbia University

9. Department of Biomedical Informatics, Columbia University

10. Department of Biostatistics, Mailman School of Public Health, Columbia University

11. Department of Environmental Health Sciences, Mailman School of Public Health, Columbia University

12. Center for Precision Medicine & Genomic/Department of Medicine and Division of Ethics, Department of Medical Humanities and Ethics, Columbia University

13. Division of General Medicine, Department of Medicine, Columbia University

#### U01HG013276

Stephanie A. Christenson^14^, Neeta Thakur^14^, John R. Balmes^15,16^, Julianna Deardorff^17^, Jocelynn Arellanes^14^, Karina J. Barragan^16^, Rosemarie de la Rosa^16^, Laura M. Diaz^16^, Jackelyn C. Ledesma^14,7^, Jacqueline León^14^, Christine Y. Lin^14^, Nolan McCormick^16^, Srilaxmi Nerella^14^, Andrew V. Nguyen^14^, Isabella Vasquez Del Mercado^16^, Morgan Ye^14^

14. Division of Pulmonary, Critical Care, Allergy, and Sleep Medicine, Department of Medicine, University of California, San Francisco

15. Department of Medicine, University of California, San Francisco, CA, USA

16. Division of Environmental Health Sciences, School of Public Health, University of California, Berkeley, CA

17. Division of Community Health Sciences and Center for Environmental Research and Community Health, School of Public Health, University of California, Berkeley

#### U01HG013288

Lida Chatzi^18^, Max T. Aung^18^, Tanya L. Alderete^19^, Lourdes Baezconde-Garbanati^18^, Brittney O. Baumert^18^, Sirikorn Boonsawat^20^, Rosalba Cain^21,7^, Tamar Galindo^18^, Jesse A. Goodrich^18^, Rohit Kohli20, Zhenjiang Li^18^, Shiwen Li^22^, Ronald Martinez^20^, Qi Meng^18^, Fotini Tania Mitsinikos^20^, Nathan R. Pavlovic^23^, Isabella F. Valenzuela^20^, Alaina P. Vidmar^24^

18. Department of Population and Public Health Sciences, University of Southern California Keck School of Medicine

19. Department of Environmental Health and Engineering, Bloomberg School of Public Health, Johns Hopkins University, Baltimore, Maryland 21205, United States

20. Division of Gastroenterology, Hepatology, and Nutrition, Children’s Hospital Los Angeles

21. Southern California Clinical and Translational Science Institute, University of Southern California, Keck School of Medicine

22. Department of Public Health Sciences, Thompson school of social work & public health, University of Hawaiʻi at Mānoa

23. Sonoma Technology, Inc.

24. Division of Endocrinology, Children’s Hospital Los Angeles

#### U01HG013189

Louise C. Laurent^25^, Marni B. Jacobs^25^, Kathryn J. Lindley^26^, Ravi V. Shah^6^, Sierra Adkins^25^, Jennifer E. Below^3,27^, Anesha M. Byrd^28^, Anelizze Castro-Martinez^25^, Mariko Horii^29^, Kayle D. Karcher^25,7^, Brianna L. Rodriguez^25^, Maya L. Selvaraj^25^, Valentina Stanley^25^, Carolina Thorlund-Diaz^25^, Beatriz Valeiro^25^

25. Department of Obstetrics, Gynecology, and Reproductive Sciences, University of California San Diego, La Jolla, CA

26. Cardiovascular Division, Vanderbilt University Medical Center, Nashville, TN

27. Department of Medicine, Division of Genetic Medicine and Clinical Pharmacology, Vanderbilt University Medical Center, USA;

28. Department of Cardiovascular Medicine, Vanderbilt University Medical Center, Nashville, TN

29. Department of Pathology, University of California San Diego, La Jolla, CA

#### U01HG013275

Tanika N. Kelly^30^, Maria Argos^31^, Ana C. Ricardo^32^, Hua Yun Chen^33^, Martha Daviglus^34^, Robert Sargis^30^, Tamar Sofer^35^, Diana Alvarado^30^, Eryn Brazil^30^, Monica Duron^30^, Renetta Glass Ward^7^, Zhijie Huang^30^, Brenna Kirkpatrick^30^, James P. Lash^36^, Brian Layden^30^, Xiang Li^30^, Anne Muldoon^31^, Yingyue Ni^31^, Yang Pan^30^, Eve Pinsker^37^, Margaret Quaid^31^, Habiba Sultana^30^, Xiao Sun^30^, Yixi Sun^33,38^, Pei Wen Tung^31^

30. Division of Endocrinology, Diabetes, and Metabolism, University of Illinois Chicago, Chicago, IL

31. Department of Environmental Health, Boston University School of Public Health, Boston, MA

32. Department of Medicine, Division of Nephrology, University of Illinois Chicago, Chicago, IL

33. Division of Epidemiology and Biostatistics, School of Public Health, University of Illinois Chicago, Chicago, IL, USA

34. Institute for Minority Health Research, University of Illinois Chicago

35. Cardiovascular Division, Beth Israel Deaconess Medical Center, Boston, MA 02461

36. Division of Nephrology, University of Illinois Chicago, Chicago, IL 60612

37. Community Health Sciences, School of Public Health, University of Illinois at Chicago

38. Division of Endocrinology, Diabetes and Metabolism, Department of Medicine, College of Medicine, University of Illinois Chicago, Chicago, IL, USA

### Omics Production Center Award (contact PI, MPI, Co-I, other members, all alphabetical by last name)

#### U01HG013227

Gary J. Patti^39,40^, Ting Wang^41^, Leah P. Shriver^39,40^, Derek Albracht^42^, Elizabeth L. Appelbaum^42^, Edward A. Belter, Jr^42^, Ju-Fang Chang^41^, Lucy Chen^41^, Shihua Dong^41^, Robert S. Fulton^41,42^, Byoungsook Goh^39,40^, Triston Groff^39,40^, Michael Heinz^42^, Jihyun Kang^39,40^, Cassandra G. Kempf^39,40^, Amy Li^41^, Daofeng Li^41^, Ping Liu^43^, Juan F. Macias-Velasco^41^, Yiling Mi^39,40^, Heather Schmidt^41^, Michaela Schwaiger-Haber^39,40^, Chad Tomlinson^42^, Nikki L. Trapp^42^, Xiaoyun Xing^41^

39. Departments of Chemistry and Medicine, Washington University, St. Louis, MO

40. Center for Mass Spectrometry and Metabolic Tracing, Washington University, St. Louis, MO

41. Department of Genetics, Washington University School of Medicine, St. Louis, MO, USA

42. McDonnell Genome Institute, Washington University School of Medicine, St. Louis, MO, USA

43. GTAC@MGI, Washington University in St. Louis

### Data Analysis and Coordination Center Award (contact PI, MPIs, Co-Is, other members, all alphabetical by last name)

#### U01HG013198

Zhiping Weng^44^, Anshul Kundaje^45,46^, William S. Noble^47^, Nezar Abdennur^44^, Honghuang Lin^48^, Jill E. Moore^44^, Gregory Andrews^44^, Jonathan Fisher^44^, Vivian Hecht^45^, Mansi Khandpekar^44^, Matthew C. Lacadie^44^, Alyssa Lima^44^, Jair Meza^44^, Ananthan Nambiar^47^, Nishigandha Phalke^44^, Huong Phan^44^, Christian Ramirez^44^, Nicole Shedd^44^, Ross W. Swofford^44^, Vedat Yilmaz^44^, Doudou Yu^47^

44. Department of Genomics and Computational Biology, UMass Chan Medical School

45. Department of Genetics, Stanford University, Stanford, CA, USA

46. Department of Computer Science, Stanford University, Stanford, CA, USA

47. Department of Genome Sciences, University of Washington

48. Department of Medicine, UMass Chan Medical School

### NIH Program Management

#### National Cancer Institute

Melissa Rotunno

## Supplementary Figures

**Supplementary Figure 1.**
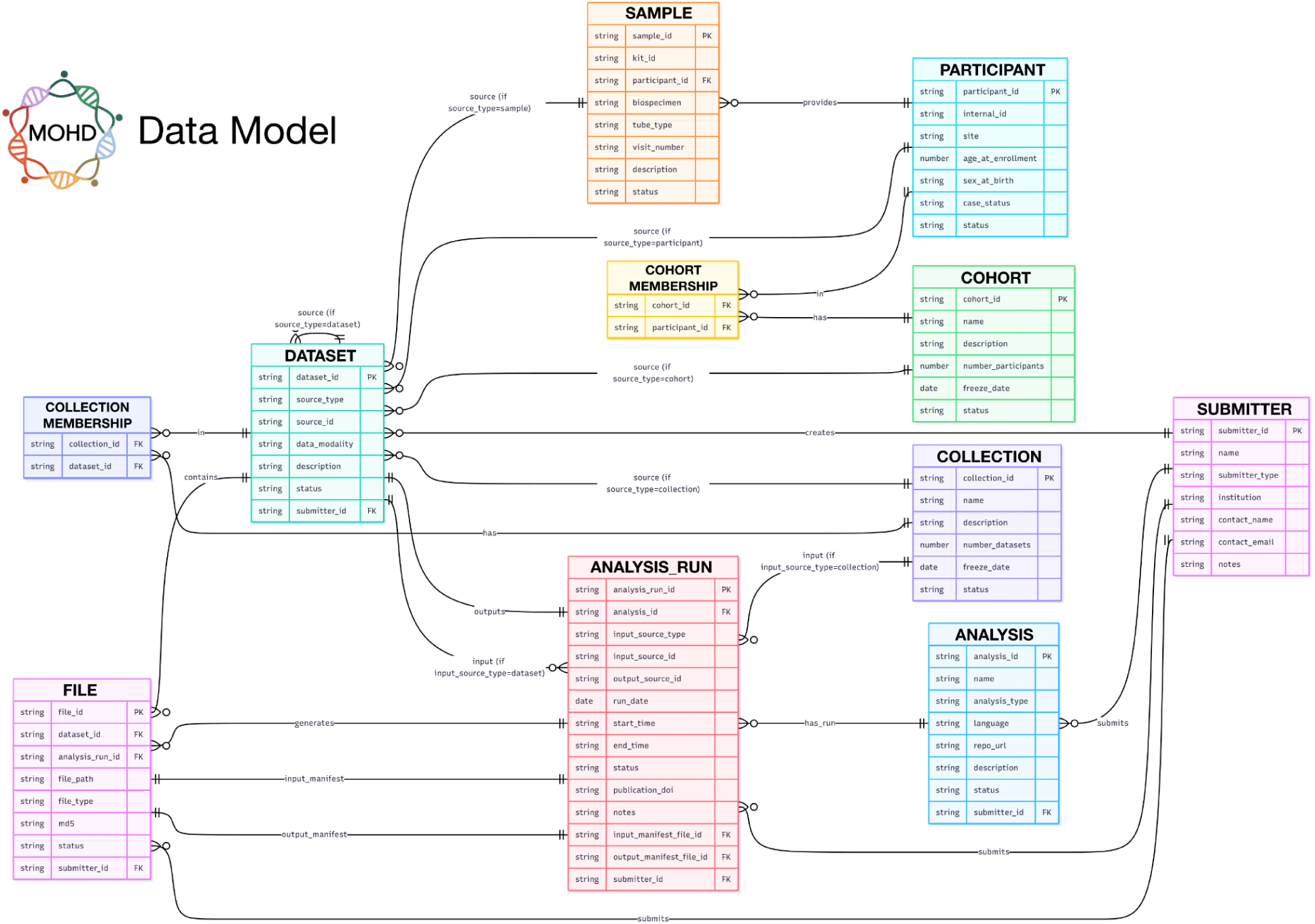
Detailed schema of the MOHD data model. Entity-relationship diagram of the MOHD data model showing core entities, attributes, and their relationships. Primary entities include participants, samples, datasets, collections, cohorts, analyses, analysis runs, files, and submitters. Datasets serve as the central entity and are linked to their source (participant, sample, dataset, or collection) and associated files. Analysis and analysis run entities capture computational provenance, including defined workflows and their specific executions, with explicit tracking of inputs and outputs. Membership tables (cohort membership and collection membership) define many-to-many relationships between participants and cohorts, and between datasets and collections. Together, these entities and relationships enable consistent tracking of longitudinal data generation, processing, and reuse within the MOHD framework.

